# Short-term evolution under copper stress increases probability of plasmid uptake

**DOI:** 10.1101/610873

**Authors:** Uli Klümper, Arnaud Maillard, Elze Hesse, Florian Bayer, Stineke van Houte, Ben Longdon, Will Gaze, Angus Buckling

**Author notes:** corresponding author: Uli Klümper, College of Life and Environmental Sciences, ESI Building, University of Exeter, TR109FE Penryn, Cornwall, United Kingdom, ORCID: 0000-0002-4169-6548.

## Abstract

Understanding plasmid transfer dynamics remains a key knowledge gap in the mitigation of antibiotic resistance gene spread. Direct effects of exposure to stressors on plasmid uptake are well monitored. However, it remains untested whether evolution of strains under stress conditions modulates subsequent plasmid uptake. Here, we evolved a compost derived microbial community for six weeks under copper stress and non-exposed control conditions. We then tested the ability of isolated clones from both treatments to take up the broad host range plasmid pKJK5 from an *E.coli* donor strain. Clones pre-adapted to copper displayed a significantly increased probability to be permissive towards the plasmid compared to those isolated from the control treatment. Further, increased phylogenetic distance to the donor strain was significantly and negatively correlated with plasmid uptake probabilities across both treatments.

## Text

Conjugal plasmids serve as main means of bacterial evolutionary adaptation to environmental stressors (Norman et al., 2009). The spread of plasmids encoding antibiotic resistance is considered a major threat to human health (WHO, 2014). Crucially, understanding plasmid spread dynamics remains a key knowledge gap (Smalla et al., 2018). Direct exposure to environmental stressors such as antibiotics (Slager et al., 2014), non-antibiotic pharmaceuticals (Wang et al., 2018) or metals (Klümper et al., 2017) can modulate immediate plasmid uptake in single strains and across microbial communities. This effect can either originate from direct selection or as a by-product of a general stress response. While immediate stress effects on plasmid uptake are well monitored, it remains untested whether ecological or evolutionary selection under stress conditions results in phenotypes with intrinsically higher plasmid permissiveness. There is evidence that stress or other environmental change can select for increased mutation (Pal et al., 2007) and recombination (Cooper, 2007) rate in bacteria. We therefore hypothesize that more permissive bacteria might also be favoured, as a result of horizontally acquired adaptive stress resistance.

Here, we tested if evolution in a microbial community exposed to metal stress has an effect on the plasmid uptake ability of individual clones. To infer a causal relationship between exposure to copper and subsequent plasmid uptake, we set up experimental compost communities in sterile compost following the protocol of Hesse et al. (2018). Hence, all treatments started off with the same community and level of permissiveness. Microcosms were incubated (75% humidity, 26°C), and twice (after 1 and 21 days) supplemented with either 2 ml filter-sterilized 0.25M CuSO_4_ or ddH_2_0. We here tested a total of 66 clones (27 copper, 39 control) that were isolated following 6 weeks of evolution. Copper tolerance increased significantly in clones pre-adapted to copper (Hesse et al., 2018). Clones were 16S sequenced using the 27F primer and a phylogenetic tree was constructed using mothur v1.41.1 (Schloss et al., 2009). Isolates belonged to 4 different phyla: Actinobacteria, Bacteroidetes, Firmicutes and three classes of Proteobacteria (α, β & γ) (Figure 1). Further, no significant difference in the distribution of isolates between metal and control treatment in the phylogenetic tree was detected (P-test, p=0.163, P-score=18).

**Fig. 1:**
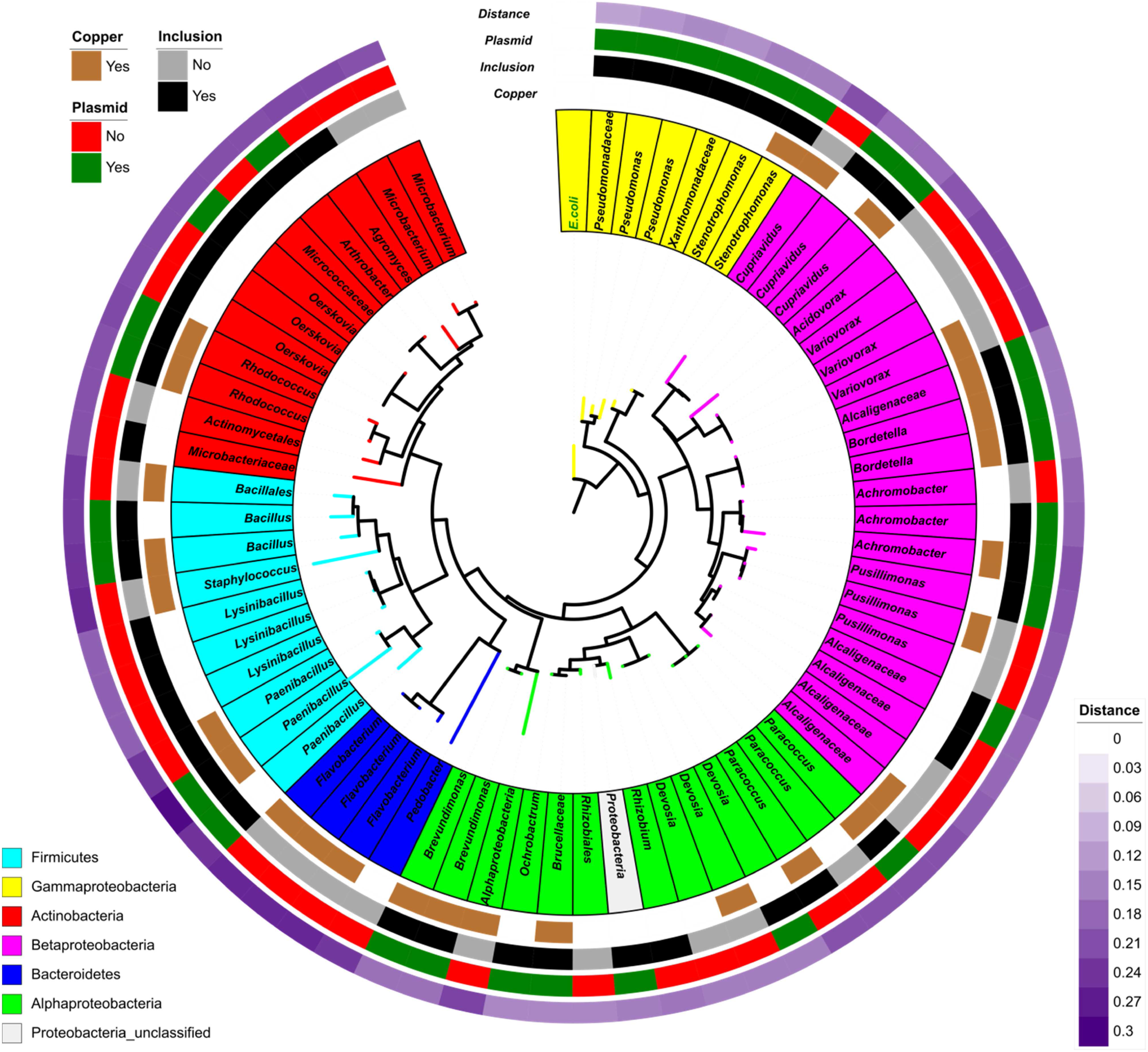
Phylogenetic tree of the 66 isolates and the donor strain *E.coli*. Clone labels are color-coded based on phylogenetic identification. The 4 heatmap rings around the tree display: A) Isolation from the copper treatment (brown) or the control treatment (white). B) Inclusion (black) and exclusion (grey) from the study based on ability to grow on citrate while displaying susceptibility to tetracycline. C) Ability to take up plasmid pKJK5 in the experimental setup (Yes = green; No = red). D) Phylogenetic distance from the *E.coli* donor strain.

To test permissiveness towards broad host range plasmid pKJK5 (Klümper et al., 2015; Li et al., 2018) each strain was mixed at 1:1 ratio with donor strain *Escherichia coli* MG1655::*Km*^*R*^*-Lpp-mCherry* hosting the IncP- 1ε conjugative plasmid pKJK5::*gfpmut3b* (Bahl et al., 2007; Klümper et al., 2014). Experiments were carried out in LB medium in absence of any selective pressure, centrifuged for 2 minutes at 10000x*g* to ensure cell-to-cell contact and incubated (24 h; 28°C). Cells were harvested, resuspended and inoculated on M9 minimal media plates supplemented with 10 mM citrate and 10 μg/mL tetracycline. Citrate as the single carbon source counter-selects against the *E.coli* donor strain, while tetracycline selects for acquisition of the tetracycline resistance encoding plasmid. Upon successful growth, green fluorescence, repressed in the donor strain but expressed upon transfer in transconjugants, was confirmed using fluorescence microscopy. Out of the 66 strains 42 were able to grow on citrate medium and displayed susceptibility to tetracycline. These were included in subsequent analysis with 71.4% permissive to plasmid pKJK5 (Figure 1). However, permissiveness differed strongly across phyla. Out of 25 proteobacterial strains, belonging to the same phylum as the *E.coli* donor, 22 (88%) took up the plasmid, while only 47% of gram positive strains (8/17) were permissive.

We subsequently fitted a logistic regression model (Figure 2) with the isolates evolutionary background and phylogenetic distance to the donor strain as explanatory variables to predict plasmid uptake probability. Both copper background (ANOVA χ2-test, p=0.0416, dF=39) and phylogenetic distance (ANOVA χ2-test, p=0.0033, dF=40) proved statistically significant in predicting plasmid uptake. However, while increasing distance to the donor strain had a negative effect on plasmid uptake probabilities, strains pre-adapted to copper were more likely to take up pKJK5 compared to non-adapted strains (Figure 2).

**Fig. 2:**
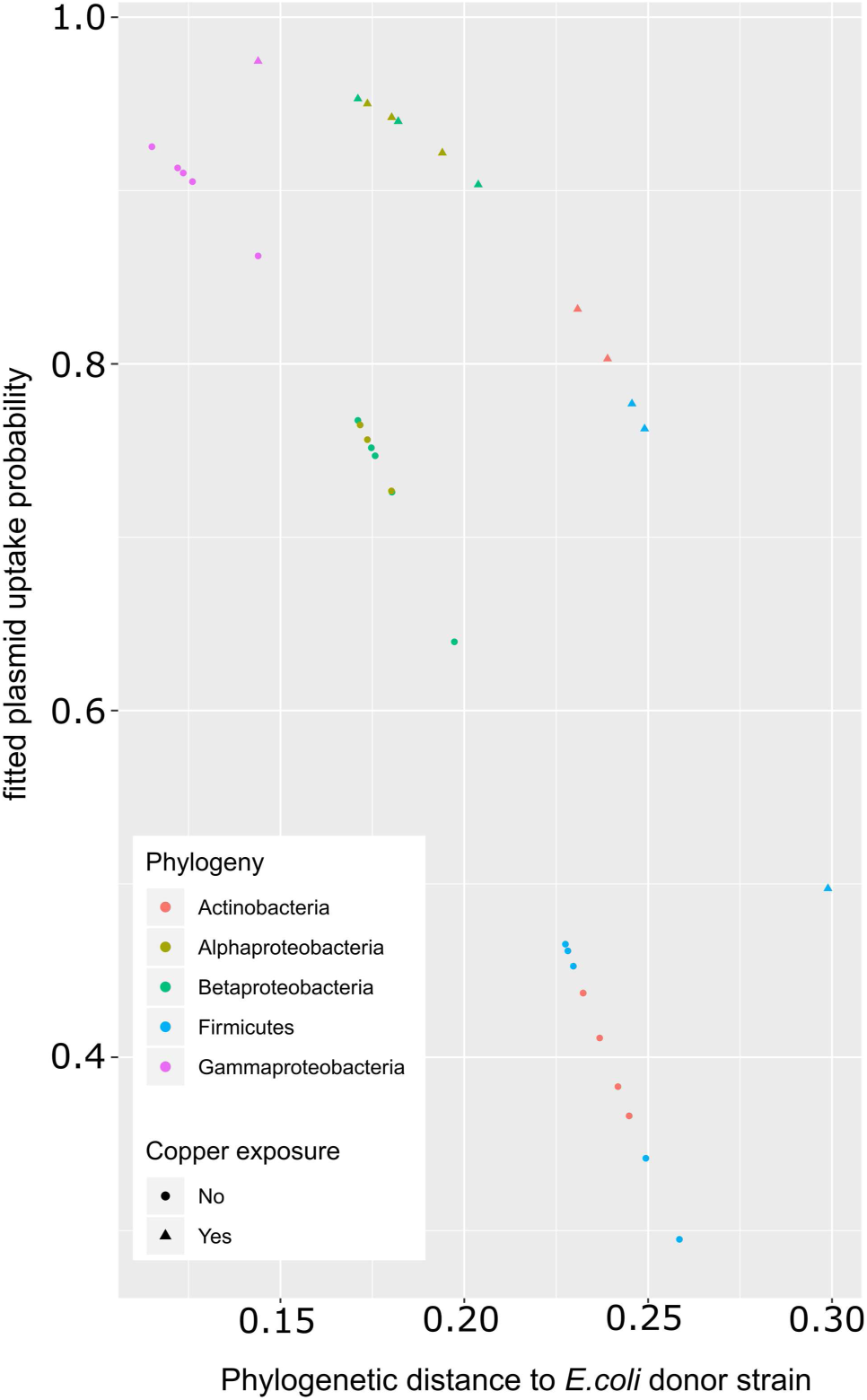
Predictive modelling of plasmid uptake probability based on phylogenetic distance to donor strain *E.coli* and previous metal exposure. Fitted values of logistic regression for the 42 included strains based on the model *p*_ij_ = *α*_*i*_ + *β* × *d* (*p*_ij_: the probability of pKJK5 plasmid uptake in our experimental setup; α_i_: the effect associated with evolution under metal stress conditions; β: weighing parameter of the distance to the donor strain *E.coli*; *d*: distance to donor strain *E.coli*). Symbols are colour coded based on phylogenetic identification. Isolates from copper exposure are shown as triangles, isolates from the control treatment as circles.

Applying strong selection, such as high concentrations of metals, can have major effects on bacterial ecology and evolution (Giller et al., 1998). Here, under copper exposure phenotypes with increased likelihood to be permissive either evolved or were positively selected for. However, these two very different underpinning mechanisms cannot be distinguished at this point, as evolved communities different in their composition as a result of ecological species sorting (Hesse et al., 2018).

Subsequent work on single tractable species will explore the exact mechanisms underlying increased permissiveness. If, as a direct consequence of stress, more highly permissive bacteria were to survive better, they should also intrinsically host more plasmids, consequentially making them less permissive due to plasmid incompatibility or entry exclusion (Garcillán-Barcia and de la Cruz, 2008). This suggests that higher permissiveness might likely be an indirect by-product due to genetic changes in for example stress response, membrane permeability or pilus expression. Further, immunity towards plasmids could be evolutionary lost; CRISPR-Cas or less specific abortive infection systems can be lost under conditions when they bear immunity to horizontally transferred, potentially beneficial genes (Jiang et al., 2013).

Therefore, the conjugative host range can be increased under metal stress. The conjugative host range is generally assumed as much broader than the persistence host range (De Gelder et al., 2007). Consequently, the genomic signature of IncP-type plasmids suggests Proteobacteria as their main evolutionary hosts (Suzuki et al., 2010), which we also found to display far higher probabilities of plasmid uptake.

In strains with a higher degree of phylogenetic distance to the donor stability of pKJK5 might be very low and thus lost within few generations. However, long-term metal stress has been proven to elevate the retention of plasmids, even for those not coding for metal resistance (Smets et al., 2003). Even though plasmid-host co-evolution might reach an epidemiological dead end in some, likely more distant strains, such spill over remains crucial for propagation of antibiotic resistance. Unique events of recombination with the chromosome could happen before plasmid loss, especially since pKJK5 like many plasmids hosts a highly recombinative integrative element (Bahl et al., 2007). An increase of plasmid uptake ability and potentially retention time under metal stress conditions would consequently increase the likelihood and extent of such recombination events and foster the spread of antibiotic resistance.

## Acknowledgments

UK received funding from the European Union’s Horizon 2020 research and innovation program under Marie Sklodowska-Curie grant agreement no. 751699. UK, AB and WG were supported through an MRC/BBSRC grant (MR/N007174/1). AM received internship funding from the “Conseil Régional d’Ile de France”.

## Author contributions

UK, AM, SvH, WG and AB conceived the study and designed experiments; AM performed permissiveness assays with support from UK; EH performed evolution experiment and strain isolation; FB performed molecular work and sequencing; UK, AM, EH, BL analysed data; UK and AB wrote the manuscript.

## Materials & Correspondence

All correspondence and material requests should be addressed to UK.

